# mTORC1 hyperactivation causes sensory integration deficits due to habenula impairment

**DOI:** 10.1101/2022.07.20.500762

**Authors:** Olga Doszyn, Magdalena Kedra, Justyna Zmorzynska

**Affiliations:** Laboratory of Molecular and Cellular Neurobiology, International Institute of Molecular and Cell Biology in Warsaw

**Author notes:** corresponding author: Justyna Zmorzynska, **Email:**. equal contribution (Olga Doszyn, Magdalena Kedra). **Author Contributions (CRediT):** OD: Investigation, Formal analysis, Writing – Review & Editing; MK: Investigation, Formal analysis; JZ: Conceptualization, Investigation, Formal analysis, Writing – Original Draft, Writing – Review & Editing, Visualization, Supervision, Project administration, Funding acquisition. **Competing Interest Statement:** The authors declare no competing interests.

**Keywords:** mTORC1 hyperactivation, habenula, Tuberous Sclerosis Complex (TSC)

## Abstract

Mechanistic target of rapamycin complex 1 (mTORC1) is an integration hub for extracellular and intracellular signals necessary for proper brain development. Hyperactivation of mTORC1 is found in many developmental diseases, including autism spectrum disorder (ASD). Atypical reactivity to sensory stimuli is often found in patients with ASD. The most frequent hereditary cause of ASD is Tuberous Sclerosis Complex (TSC), in which inactivating mutations in the *TSC1* or *TSC2* genes result in hyperactivation of the mTORC1 pathway. We have discovered that the zebrafish model of TSC, *tsc2*^*vu242/vu242*^ mutants lack light preference, a behavior requiring integration of multiple sensory inputs. Here, we show that the lack of light preference in *tsc2*^*vu242/vu242*^ zebrafish is caused by aberrant sensory integration of light stimuli in the left dorsal habenula. Single-cell calcium imaging analysis revealed that *tsc2*^*vu242/vu242*^ fish showed impaired function of the left dorsal habenula, in which neurons exhibited higher activity and lacked habituation to the light stimuli resulting in atypical response to light. Lack of light-preference behavior and abnormal neuronal activity in the left dorsal habenula were rescued by rapamycin, indicating that hyperactive mTorC1 causes aberrant habenula function and impaired sensory integration resulting in lack of light preference. Our results link sensory integration deficits seen in TSC patients suffering from ASD with hyperactive mTORC1 and suggest that mTORC1 hyperactivity contributes to atypical reactivity to sensory stimuli in ASD.

## Introduction

Mechanistic target of rapamycin complex 1 (mTORC1) is an integration hub for extracellular and intracellular signals that controls cell homeostasis by regulating translation, protein degradation, transcription and RNA processing, and cytoskeleton dynamics [1]. mTORC1 is necessary for proper brain development and coordinates proliferation, migration, differentiation, synaptogenesis, and neuronal activity in the brain [1]. Hyperactivation of mTORC1 is a hallmark of many developmental diseases, including autism spectrum disorder (ASD), which has a prevalence of 1% in general population. ASD symptoms include social deficits, atypical reactivity to sensory stimuli, repetitive behaviors, and speech delay [2–4].

Tuberous Sclerosis Complex (TSC) is an exemplary genetic disease with mTORC1 hyperactivation. Inactivating mutations in the *TSC1* or *TSC2* genes cause lack of functional TSC1-TSC2 complex and result in hyperactivation of mTORC1 pathway [1]. Patients with TSC suffer from epilepsy, benign tumors, and TSC-associated neuropsychiatric disorders (TANDs). These neuropsychiatric disorders occur in more than 90% of the TSC patients and do not fully correlate with tumor or seizure burden (reviewed in [5]). Approximately 40% of TSC patients have ASD, which makes TSC the most frequent hereditary cause of ASD [6]. However, the underlying pathomechanisms of TSC-associated ASD are still obscure.

Aberrant sensory processing leading to “sensory overload” is a hallmark of ASD [7], however, the mechanism underlying this deficit is not fully understood. Individuals with ASD often attempt to avoid visual stimulation [8] and show high levels of mTORC1 activity [3]. We investigated light processing in the light preference paradigm in the zebrafish model of TSC [9] to identify mTorC1 as an underlying cause for aberrant activity of left dorsal habenula (LdHb) and atypical response to light in the light-preference test. Our results suggest that mTORC1 hyperactivity contributes to atypical reactivity to sensory stimuli in ASD.

## Results

The light-preference assay measures anxiety by comparing times spent in the light and dark compartments. Increased preference for light of zebrafish larvae is indicative of anxiety-like behavior. We have previously shown that *tsc2*^*vu242/vu242*^ mutant zebrafish exhibit anxiety-like behavior and elevated cortisol levels [10]. However, in the light-preference assay, *tsc2*^*vu242/vu242*^ fish did not present increased light preference indicative of anxiety (Fig.1A-C). They exhibited decreased light preference compared to wild-type (wt) siblings, which strongly preferred the light compartment, although the total time moving was similar among genotypes (Fig.1E). The *tsc2*^*vu242/vu242*^ fish also did not exhibit visual problems [10]. The lack of light preference of *tsc2*^*vu242/vu242*^ was not prevented by pretreatment with anxiolytic drug ANA-12 (Fig.1B-D), which was shown before to block anxiety-like behavior in *tsc2*^*vu242/vu242*^ [10]. Thus, the lack of light preference was not associated with anxiety. The aberrant response to light in *tsc2*^*vu242/vu242*^ was also not associated with seizures as anti-epileptic vigabatrin (VGN) did not inhibit this phenotype either (Fig.1B-D). Interestingly, the pretreatment with a direct mTorC1 inhibitor rapamycin reversed the aberrant light response of *tsc2*^*vu242/vu242*^ mutants (Fig.1B,C). These results indicate that hyperactive mTorC1 underlies the lack of light-preference behavior of *tsc2*^*vu242/vu242*^ fish.

**Figure 1.**
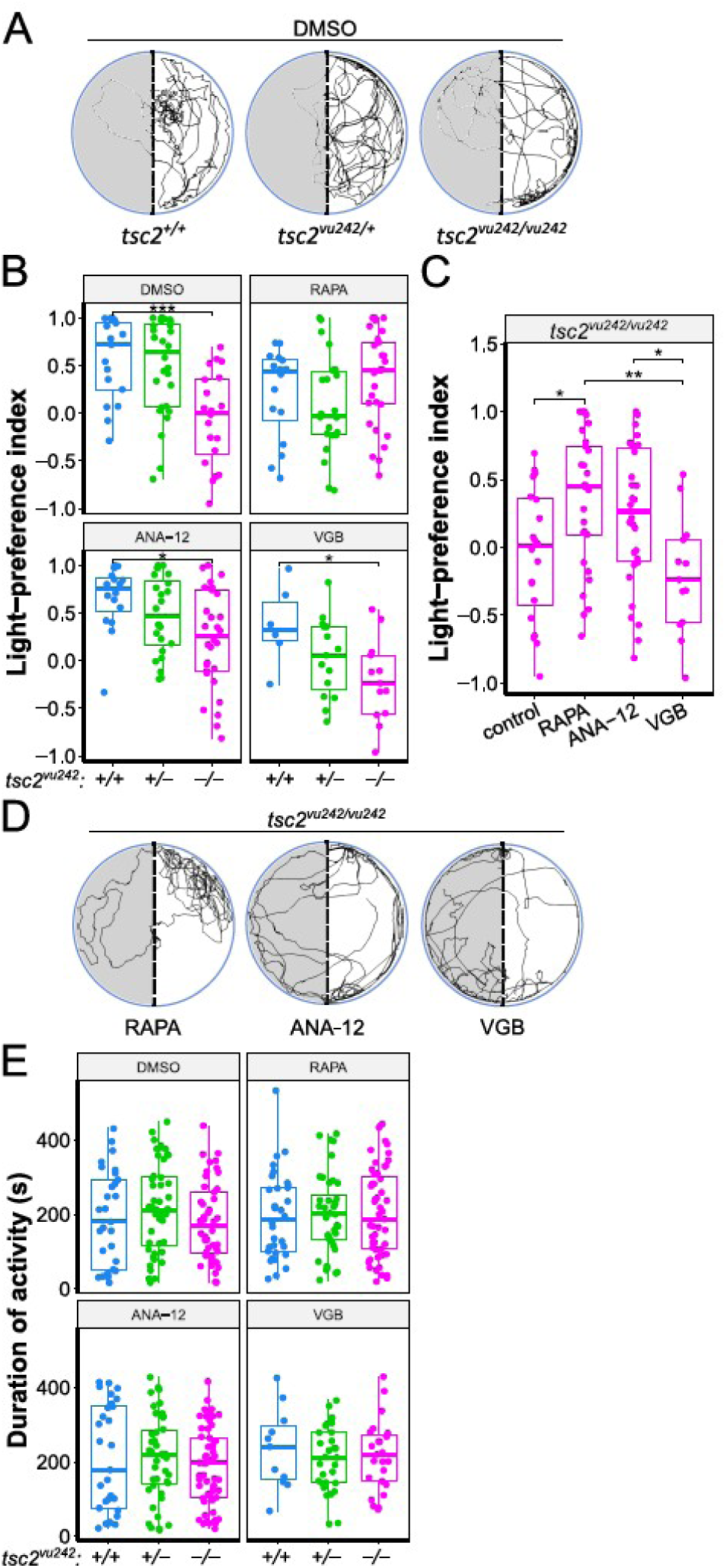
Light-preference test in *tsc2*^*vu242*^ fish. A. Exemplary tracks of *tsc2*^*vu242*^ fish from the light-preference test. B. Light-preference index in *tsc2*^*vu242*^ fish. C. Light-preference index in *tsc2*^*vu242/vu242*^ mutants with comparison statistics between treatments. D. Exemplary tracks of *tsc2*^*vu242/vu242*^ mutant treated with rapamycin (RAPA), ANA-12, or VGB from the light-preference test. E. Cumulative activity of *tsc2*^*vu242*^ fish during the light-preference test.

The lack of light preference in the *tsc2*^*vu242/vu242*^ mutants, otherwise exhibiting increased anxiety-like behaviors, can be indicative of impaired sensory processing of the light stimulus or impaired integration of sensory input and internal states which converge to an aberrant behavioral response, resulting in decreased light preference. Habenula integrates various stimuli and regulates light-preference behavior [11–13]. To confirm mTorC1 hyperactivity in the habenulae of the *tsc2*^*vu242/vu242*^, we checked phosphorylation levels of its downstream target: ribosomal protein s6 (Rps6). We found that the number of cells positive for phosphorylated Rps6 (pRps6) in the *tsc2*^*vu242/vu242*^ mutants was increased specifically in left dorsal habenula (LdHb) compared to wt siblings (Fig.2A,B). Also, the pRps6 intensity levels per cell were higher in the *tsc2*^*vu242/vu242*^ LdHb neurons than in the wt siblings (Fig.2C). The pRps6 levels were decreased by rapamycin pretreatment in both *tsc2*^*vu242/vu242*^ mutants and wt siblings (Fig.2).

**Figure 2.**
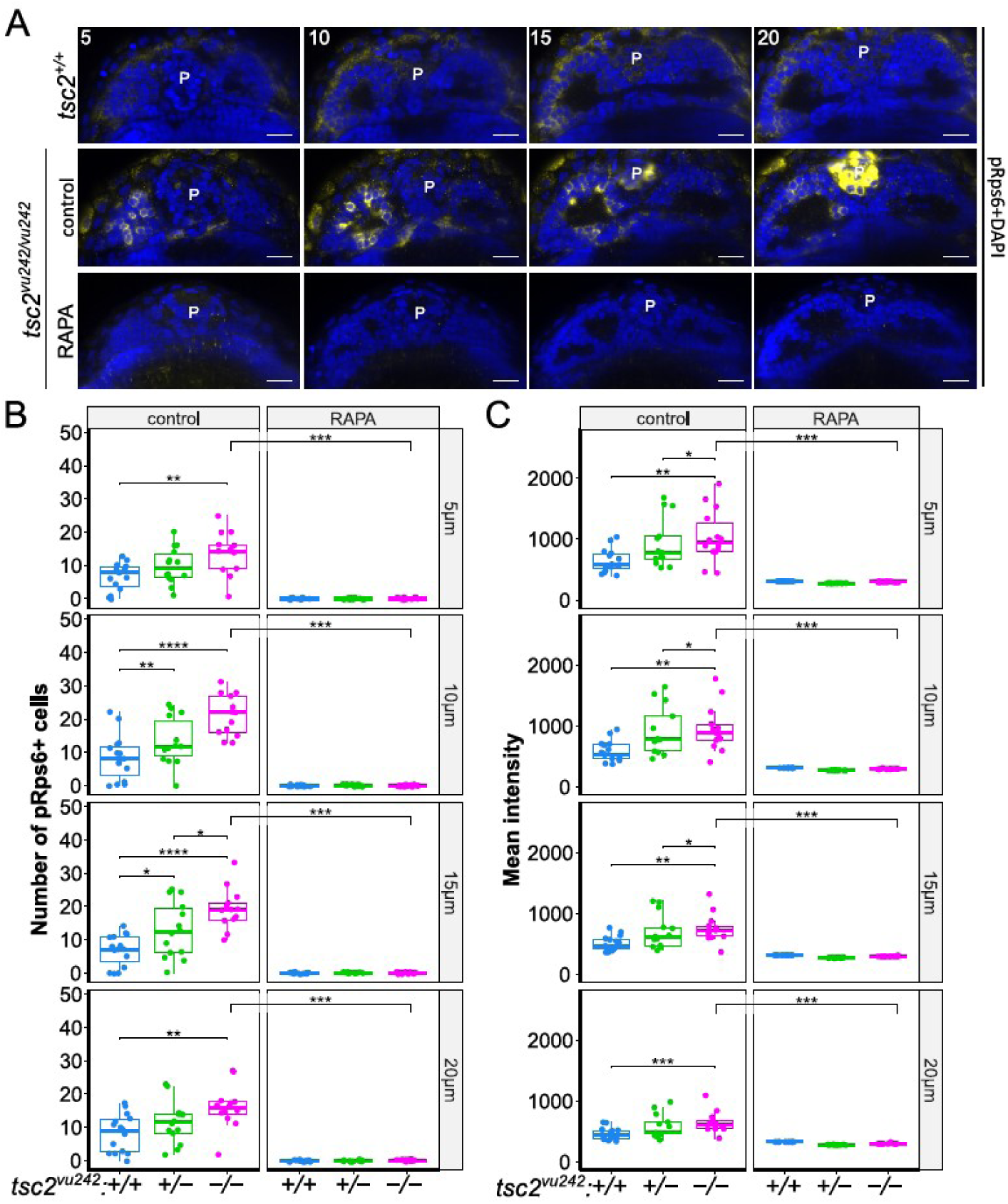
mTorC1 activation in *tsc2*^*vu242*^ fish. A. Representative optical sections through habenula of *tsc2*^*vu242*^ fish at 5, 10, 15, and 20 μm from the top. PRps6 – yellow, nuclei – blue. P – pineal complex. Scale bars, 20 μm. B. Number of pRps6-positive cells in LdHb of *tsc2*^*vu242*^ fish. C. Quantification of mean intensity of pRps6 fluorescence from LdHb of *tsc2*^*vu242*^ fish.

LdHb contains light-responsive neurons and is responsible for mediating light-preference behavior in zebrafish larvae and its impairments result in lack of light preference [11, 13]. Thus, we performed 3D time-lapse imaging of the activity of LdHb neurons in *Tg(HuC:GCaMP5G);tsc2*^*vu242*^ expressing GCaMP5G under neuron-specific promoter. The single-cell analysis revealed that neuronal activity in LdHb was increased in *tsc2*^*vu242/vu242*^ mutants compared to wt siblings (Fig.3). Neuronal activity dynamics across time revealed that after light stimulation, the activity of neurons increased in wt LdHb but decreased over time indicative of habituation to constant light stimulus. In contrast, the activity of neurons in *tsc2*^*vu242/vu242*^ LdHb was lower at the initial stimulation but increased over time (Fig.3C,D). Rapamycin pretreatment decreased neuronal activity of *tsc2*^*vu242/vu242*^ LdHb (Fig.3) implicating mTorC1 hyperactivation in the aberrant LdHb activity.

**Figure 3.**
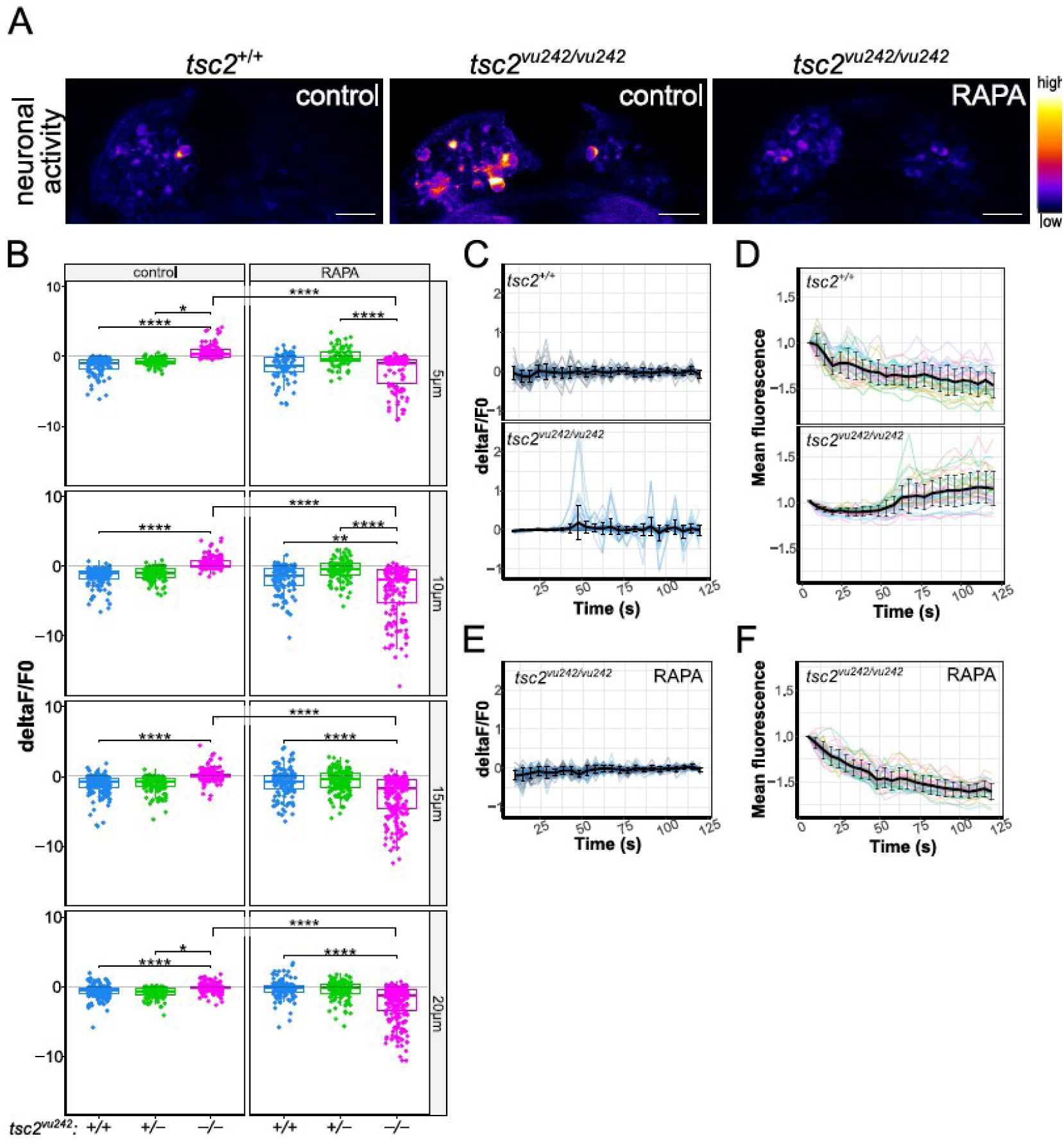
Neuronal activity in LdHb in *tsc2*^*vu242*^ fish. A. Representative images of neuronal activity in the habenulae of *tsc2*^*vu242*^ fish at 10 μm from the top. Scale bars, 20 μm. B. Cumulative activity of the *tsc2*^*vu242*^ LdHb. C. Neuronal activity change over time in the *tsc2*^*vu242*^ LdHb at 10 μm from the top. D. Normalized mean GCaMP fluorescence over time in the *tsc2*^*vu242*^ LdHb at 10 μm from the top. E. Neuronal activity change over time in the *tsc2*^*vu242/vu242*^ mutant’s LdHb at 10 μm from the top after RAPA pretreatment. F. Normalized mean GCaMP fluorescence over time in the *tsc2*^*vu242/vu242*^ mutant’s LdHb at 10 μm from the top after RAPA. C-F. The mean with SD is shown in black.

The left habenula receives afferent inputs from the eminentia thalami (EmT), the pallium through stria medullaris (SM), and from the right habenula through the habenula commissure (HC) [11]. Therefore, we checked the development of these afferents in *tsc2*^*vu242/vu242*^ mutants as its alterations could facilitate impaired LdHb function. The EmT fibers that innervate LdHb are calretinin-positive, thus, we checked anti-calretinin immunofluorescence in the LdHb in whole-mount brain preparations. We determined that the mean intensity of the anti-calretinin signal was similar across genotypes (Fig.4A), suggesting proper innervation of LdHb by EmT in the *tsc2*^*vu242/vu242*^. The immunofluorescence staining against acetylated Tubulin (AcTub) of whole-mount brain preparations revealed that the lateral input to habenulae through SM was not significantly changed in *tsc2*^*vu242/vu242*^ compared to wt (Fig.4B). However, the HC was thinner in *tsc2*^*vu242/vu242*^ compared to wt controls, and pretreatment with rapamycin reversed its width (Fig.4C,D) suggesting that right habenula input to LdHb may be impaired.

**Figure 4.**
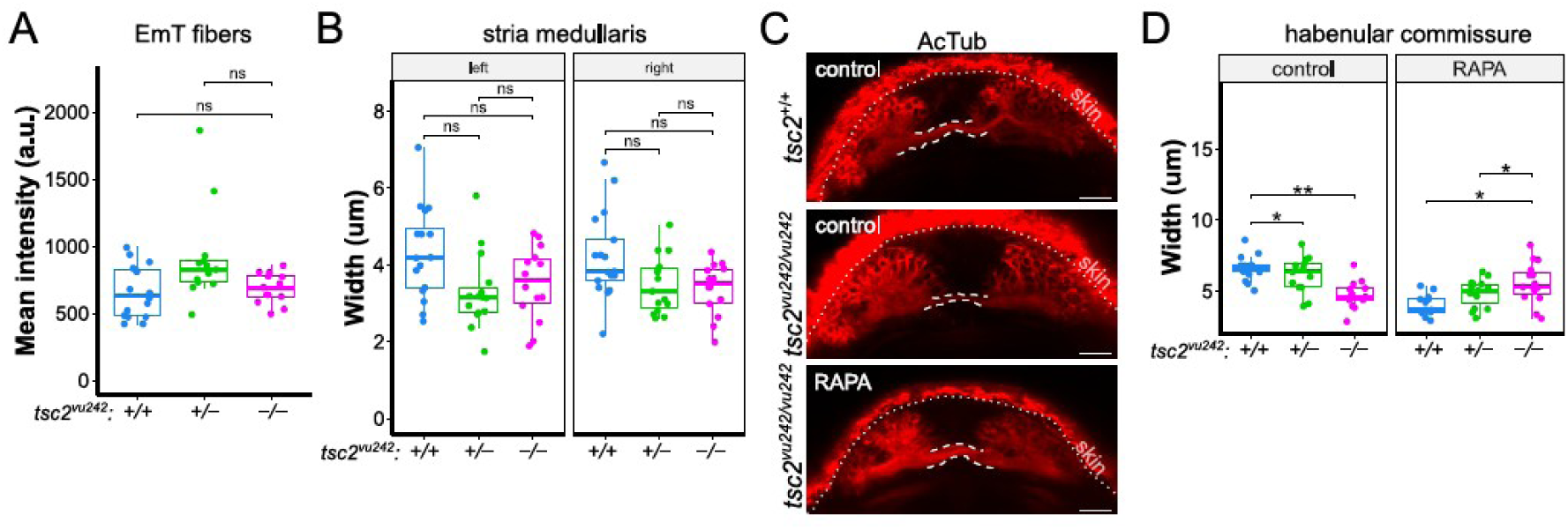
Afferent connectivity of LdHb in *tsc2*^*vu242*^ fish. A. Mean intensity of calretinin fluorescence in the left habenulae of *tsc2*^*vu242*^ fish. B. Width of stria medullaris in *tsc2*^*vu242*^ fish. C. Representative horizontal optical sections through HC (outlined) of *tsc2*^*vu242*^ fish (projection of 8 z-stacks). Scale bars, 20 μm. D. HC width of *tsc2*^*vu242*^ fish.

## Discussion

We have shown that aberrant activity of LdHb neurons correlated with hyperactivation of the mTorC1 pathway and decreased light preference in the *tsc2*^*vu242/vu242*^ mutants. The involvement of mTORC1 in neuronal activity is well documented and hyperactive mTORC1 consistently produces neuronal hyperexcitability and seizures [14]. The increased neuronal activity of LdHb neurons in *tsc2*^*vu242/vu242*^ can be indicative of decreased activation threshold which is seen in the pallium of *tsc2*^*vu242/vu242*^ and is responsible for seizures [10]. However, anti-epileptic VGB did not rescue light-preference behavior and rapamycin did reverse both, increased neuronal activity of LdHb neurons and light-preference behavior in *tsc2*^*vu242/vu242*^ fish, suggesting that LdHb activity is not induced by seizures. Instead, hyperactive mTorC1 causes aberrant LdHb function and impairs sensory integration resulting in lack of light preference in *tsc2*^*vu242/vu242*^ fish. LdHb integrates light stimuli from EmT and the right habenula with other inputs to produce light-preference behavior. In older zebrafish larvae, deactivation of LdHb by botulinum toxin decreased light preference, and activation of LdHb by optogenetic approach resulted in a preference for light in wt zebrafish [13]. In *tsc2*^*vu242/vu242*^ fish, however, the LdHb activity is impaired – low at initial stimulation, but increasing in time. It suggests that the threshold for activation may be higher but results in higher neuronal activity when crossed or that the habituation to the light stimulus is impaired in *tsc2*^*vu242/vu242*^. It is possible that intracellular signaling pathways are abnormally functioning due to hyperactive mTorC1 and therefore the synaptic inputs to LdHb are not integrated properly or timely. Our results link sensory integration deficits with hyperactive mTORC1 and suggest that rapamycin derivatives can be used to prevent atypical reactivity to sensory stimuli in ASD.

## Materials and Methods

### Zebrafish breeding, genotyping, and behavior

The following zebrafish lines were used: *tsc2*^*vu242/+*^ [9] and *tsc2*^*vu242/+*^*;Tg(HuC:GCaMP5G)* [10, 15]. Adult and larval zebrafish were bred according to international standards. The larvae were genotyped as previously described [10]. Offspring of at least two parental pairs were used in each experiment. Light-dark box test was performed as previously [10]. Lack of movement of *tsc2*^*vu242/vu242*^ mutants was mapped to non-motor seizures before [10], thus, not moving fish were excluded from the analysis. The light preference index was calculated as cumulative time spent in the dark compartment subtracted from cumulative time spent in the light compartment and divided by the total time of movement ((L-D)/(L+D)) [13].

### Additional Materials and Methods

Additional information about drug treatments, brief protocols for methods, and statistical analysis can be found in SI Appendix.

### Material and Data Availability

The manuscript and SI Appendix contain all data with the representative images to evaluate the conclusions and to reproduce the analysis. The fish lines and materials are available publicly or from the corresponding author. The single-cell calcium imaging analysis data will be made available on Github upon publication.

## Acknowledgments

We thank Kevin Ess (Vanderbilt University) for *tsc2*^*vu242/+*^ and Michael Orger (Champalimaud Foundation) for *Tg(HuC:GCaMP5G)*, the IIMCB ZCF for assistance with the adult fish, Tomasz Wegierski for Lightsheet Z.1 maintenance, and Angelika Jocek for administrative help. This work was supported by OPUS grant no. 2020/37/B/NZ3/02345 (OD, JZ) and ETIUDA grant no. 2020/36/T/NZ3/00132 (MK), both from National Science Centre, Poland.

