## Supplementary material for "mTORC1 hyperactivation causes sensory integration deficits due to habenula impairment": SI Appendix

**Extended methods**

*Drug treatments*
To prepare stock solutions, drugs were dissolved in E3 or dimethylsulfoxide and were further diluted in E3. Drugs were administered directly into E3 with the same number of dechorionated fish. Treatments included: 200 nM rapamycin (MBL International) from 2 days post-fertilization (dpf), 50 nM ANA-12 or 60 µM VGN (both from Sigma-Aldrich) 24 hours before the behavioral test.

*Whole-mount immunofluorescence*
Heads of zebrafish larvae were fixed in 4% paraformaldehyde at 4 dpf. Samples were bleached, permeabilized, blocked with 3% bovine serum albumin, and 10% goat serum, incubated with primary antibodies (anti-pRps6 antibody (Ser235/236), 1:200, CS4858, Cell Signaling Technology; anti-calretinin, 1:400, CR7697, Swant; anti-acetyl-Lys40-tubulin, 1:100, GTX16292, Genetex) at room temperature, washed, incubated with secondary antibodies (goat anti-mouse Alexa Fluor 568, 1:1000 or goat anti-rabbit, Alexa Fluor 488, 1:1000, Thermo Fisher Scientific), and mounted.

*Imaging and image analyses*
Images were acquired using a Zeiss Lightsheet Z.1 microscope (40× water immersion objective, NA = 1.3) at 1024 × 1024 pixel-resolution. Z-stacks of the images were taken with an interval of 0.5 µm for fixed samples and 1 µm for live imaging. The 3D time-lapse images were recorded every 5s for 2 min. at 28.5°C at 4 dpf. Time-lapse images included both habenulae and had an approximate range of 100 µm. Image analyses were performed using Fiji software. For pRps6, AcTub, and calretinin measurements, regions of interest (ROIs) were drawn manually and cell size, fluorescence intensity, or fiber width were measured using the measurement tool. For neuronal activity, drift correction was performed using the Tischer script, ROIs were applied manually, and then the fluorescence intensity and derivative (deltaF/F0) were calculated using the measurement tool in Fiji and Rstudio (cran.r-project.org; rstudio.com) in a semi-automated manner. All LdHb neurons from 7 fish per genotype per treatment were included in the analysis.

*Statistical analysis*
No predetermination of sample sizes was performed because the number of each genotype could not be predicted due to the random distribution and early lethality of homozygotes. Equality of variance and normality of residuals were assessed using Levene's test or Shapiro-Wilk test, respectively. When these assumptions were met, data were analyzed by either one-way or two-way analysis of variance (ANOVA) depending on number of factors with Student's t-test to correct for multiple comparisons. Otherwise, data were analyzed by Kruskal-Wallis test with *post-hoc* Wilcoxon test. The adjusted $p < 0.05$ was considered statistically significant. Data are presented as medians using boxplots with dots representing data points. Adjusted p-values are reported on figures with * for $p < 0.05$, ** for $p < 0.01$, *** for $p < 0.001$, and **** for $p < 0.0001$. In most cases, randomization was assured by lack of knowledge about the genotypes while performing the experiments. All data were analyzed with Rstudio.
